# Comparison of phylogenetic placements to pairwise alignments for taxonomic assignment of ciliate OTUs

**DOI:** 10.1101/2022.11.11.516104

**Authors:** Isabelle Ewers, Lubomír Rajter, Lucas Czech, Frédéric Mahé, Alexandros Stamatakis, Micah Dunthorn

**Affiliations:** Eukaryotic Microbiology, Faculty of Biology, University of Duisburg-Essen, Essen, Germany; Phycology, Faculty of Biology, University of Duisburg-Essen, Essen, Germany; Department of Plant Biology, Carnegie Institution for Science, Stanford, CA, United States of America; CIRAD, UMR PHIM, Montpellier, France; PHIM Plant Health Institute, University of Montpellier, CIRAD, INRAE, Institut Agro, IRD, Montpellier, France; Computational Molecular Evolution Group, Heidelberg Institute for Theoretical Studies, Heidelberg, Germany; Institute for Theoretical Informatics, Karlsruhe Institute of Technology, Karlsruhe, Germany; Natural History Museum, University of Oslo, Oslo, Norway

**Keywords:** Colpodea, diversity, environmental sequencing, phylogenetics, taxonomy

## Abstract

Taxonomic assignment of OTUs is an important bioinformatics step in analyzing environmental sequencing data. Pairwise-alignment and phylogenetic-placement methods represent two alternative approaches to taxonomic assignments, but their results can differ. Here we used available colpodean ciliate OTUs from forest soils to compare the taxonomic assignments of VSEARCH (which performs pairwise alignments) and EPA-ng (which performs phylogenetic placements). We showed that when there are differences in taxonomic assignments between pairwise alignments and phylogenetic placements at the subtaxon level, there is a low pairwise similarity of the OTUs to the reference database. We then showcase how the output of EPA-ng can be further evaluated using GAPPA to assess the taxonomic assignments when there exist multiple equally likely placements of an OTU, by taking into account the sum over the likelihood weights of the OUT placements within a subtaxon, and the branch distances between equally likely placement locations. We also inferred evolutionary and ecological characteristics of the colpodean OTUs using their placements within subtaxa. This study demonstrates how to fully analyse the output of EPA-ng, by using GAPPA in conjunction with knowledge of the taxonomic diversity of the clade of interest.

## INTRODUCTION

In environmental metabarcoding studies of protists and other microbial organisms, an important bioinformatics analysis step is the taxonomic assignment of operational taxonomic units (OTUs) (Deiner et al., 2017; Santoferrara et al., 2020). These taxonomic assignments allow for improved biological interpretations of the molecular data, such as which species or higher taxa are present and what are their functional roles in the ecosystem from which they were sampled (e.g., Beermann et al., 2021; Giner et al., 2020; ter Schure et al., 2020). Local or global pairwise alignment approaches (e.g., Needleman & Wunsch, 1970; Smith & Waterman, 1981) using a taxonomic reference database (e.g., Guillou et al., 2013; Ratnasingham & Hebert, 2007) are routinely used for such assignments in numerous metabarcoding studies. However, phylogenetic placement approaches for taxonomic assignments are being increasingly employed (Bass et al., 2018; Czech et al., 2022; Gottschling et al., 2021; Jamy et al., 2020). These two alternative approaches to assigning OTUs are beginning to be compared to each other (Hleap et al., 2021; Jamy et al., 2020). One exemplar group of organisms where we can make additional comparisons of these different approaches to taxonomic assignments are the colpodean ciliates (Dunthorn et al., 2014; Rajter et al., 2021).

Colpodean ciliates form a major ciliate clade and are frequently found in soils (Dunthorn et al., 2008; Foissner, 1993; Geisen et al., 2018; Mahé et al., 2017; Venter et al., 2018), although some also live in aquatic environments (Dunthorn et al., 2009; Dunthorn et al., 2014; Quintela-Alonso et al., 2011). Most colpodeans are bacterivorous, while a few are fungivorous or protistan predators (Adl et al., 2019; Rajter et al., 2021). Their resting cysts and r-selected reproductive strategies allow colpodean ciliates to survive and adapt to rapidly changing environmental conditions (Foissner, 1993), although sex is often not observed (Dunthorn et al., 2017). The more than 200 described colpodeans are taxonomically divided into four phylogenetically supported subtaxa: Bursariomorphida, Colpodida, Cyrtolophosidida, and Platyophryida (Adl et al., 2019; Foissner et al., 2011). For the colpodean ciliates, Rajter et al. (2021) recently released a reference alignment and corresponding phylogeny for use in phylogenetic placement analyses.

Numerous potential colpodean OTUs were found in the soils of three lowland Neotropical rainforests in an environmental metabarcoding study by Mahé et al. (2017). Using general eukaryotic primers, they sequenced the hypervariable V4 region of the small subunit ribosomal RNA SSU-rRNA locus. The V4 region in ciliates has a relatively strong phylogenetic signal compared to other short hypervariable regions (Dunthorn et al., 2014). For the taxonomic assignment of these colpodean OTUs, Mahé et al. (2017) used a global pairwise alignment approach with the Needleman-Wunsch algorithm as implemented in VSEARCH (Rognes et al., 2016) along with the Protist Ribosomal Reference (PR^2^) database (Guillou et al., 2013). To perform the pairwise alignments, VSEARCH first uses a common k-mer counter to order the reference sequences by decreasing estimated similarity with the query; then the exact similarity values are computed using a global pairwise alignment algorithm, recording only the top similarity values. VSEARCH assigns query sequences that have an identical top similarity value with more than one reference sequence are assigned by VSEARCH to the most recent common ancestor of these reference sequences. Mahé et al. (2017) retained all of the OTUs that had ≥50% similarity to a reference in the PR^2^ database, even though OTUs are normally kept only if they are ≥80% similar to a reference (Stoeck et al., 2010). The argument that Mahé et al. (2017) used for retaining OTUs with lower similarities was that the reference databases are mostly composed of sequences from marine and temperate environments, while the soil sampling for the metabarcoding was conducted in Neotropical rainforests.

Prior to the pairwise alignments, Mahé et al. (2017) implemented a phylogenetic placement data cleaning step to retain deeply divergent OTUs. The metabarcoding sequences were first aligned against the reference alignment using PaPaRa (Berger & Stamatakis, 2011), and then phylogenetically placed onto a reference tree using EPA (Berger et al., 2011). They found that for some protistan data there were some conflicts between the taxonomic assignments from the pairwise alignments and those from phylogenetic placements for some protistan taxa. Mahé et al. (2017) posited that these differences in taxonomic assignments arose in OTUs with low pairwise similarity to the taxonomic reference database, but they did not evaluate this assumption; a similar argument was made in Berger et al. (2011). In another metabarcoding study, Jamy et al. (2020) also found differences in taxonomic assignments between pairwise alignment with those from phylogenetic placements when there were low pairwise similarities to the taxonomic reference database; however, their phylogenetic assignments were derived by combining the output of phylogenetic placements with the taxonomy propagated from the nearest neighbour from a comprehensive, fully bifurcating, phylogenetic tree including the references and the queries.

Here we evaluated how taxonomic assignments inferred with phylogenetic placements compare to assignments with pairwise alignments using the colpodean OTUs from Mahé et al. (2017). Taxonomic assignments were first performed with VSEARCH. These VSEARCH-assigned colpodean OTUs were then phylogenetically placed with EPA-ng (Barbera et al., 2019) using the default options. We tested the hypothesis that differences in taxonomic assignments between pairwise alignments and phylogenetic placements occur with OTUs that exhibit a low pairwise similarity to the reference database. We then showcased how GAPPA (Czech et al., 2020) can be effectively used to evaluate cases where OTUs have more than one likely phylogenetic placement location, by evaluating their additive likelihoods in a subtaxon and their average expected distance between placement locations (EDPL, Matsen et al., 2010). We also made biological and ecological interpretations of the phylogenetically placed OTUs. The analytical approach we established here and the interpretations of the phylogenetic placements, as well as the open-source scripts to evaluate the placements across a tree, can be applied to environmental metabarcoding data of other microbial and macrobial taxa.

## METHODS

### Data and pairwise alignments

All the codes used in this study are available in an HTML file (**File S1**). Unless otherwise noted, results were visualized using ggplot2 v.3.3.2 (Wickham, 2016) and the tidyverse package v.1.3.0 (Wickham et al., 2019) in Rstudio v.4.0.3 (R Core Team, 2020).

We used 326 OTUs (**File S2**) that we obtained from the environmental metabarcoding study of three Neotropical rainforests in Mahé et al. (2017). Briefly: they extracted DNA from soil samples using general primers for the hypervariable V4 region of the SSU-rRNA locus, which was sequenced with Illumina MiSeq. These data are available at ENA/SRA under the BioProject accession number PRJNA317860. Here we recleaned and clustered the data with more updated versions of the programs. Starting with the raw sequencing data, we re-cleaned the sequences with VSEARCH v2.14.2 (Rognes et al., 2016), re-clustered into OTUs with swarm v3.0 (Mahé et al., 2022) and cleaned them again using MUMU v0.1 (https://github.com/frederic-mahe/mumu) which is a C++ implementation of LULU (Frøslev et al., 2017) for post-clustering curation of the metabarcoding data.

The centroid of each OTU, which is usually the most abundant amplicon, served as the query sequence for taxonomic assignment. For pairwise alignments, query sequences were taxonomically assigned using VSEARCH v2.14.2, with the EukRibo database v2020-10-2, which is a manually curated and extended subset of the PR^2^ database v4.12.0 (Guillou et al., 2013).

### Phylogenetic placements

For phylogenetic placement, we used EPA-ng v.0.3.8 (Barbera et al., 2019). The colpodean-specific reference alignment and reference tree came from Rajter et al. (2021), which contains full-length SSU-rRNA sequences of 62 colpodean ciliates and twelve outgroup sequences from each major ciliate group. In this alignment, ambiguous nucleotide positions are masked, except for the hypervariable V4 region. The query sequences were then aligned to the reference alignment using PaPaRa v.2.5 (Berger & Stamatakis, 2011). The main output from EPA-ng is a *jplace* file (Matsen et al., 2012), which by default contains the likelihood weight ratios for the top seven placements for each query (if there is more than one likely placement location); we also adjusted this output parameter to report the top 70 and the top 700 placements (these results, which are similar to the top 7 placements, are available in **Figure S1**). Although not evaluated here, EPA-ng does have an option *filter-acc-lwr* that tells the program to output as many placement locations as needed to reach an accumulated likelihood weight ratio value of a given total probability (e.g., 0.99). The likelihood weight ratios represent the probability to which extent an individual placement into a specific branch is supported by the data. For a given placed sequence, they sum to 1 (total probability) across all branches of the tree. In other words, the higher the likelihood weight ratio value of a specific placement on a branch of the reference tree, the more likely it is. Visualization of the distribution of the placements for each query sequence across the reference tree was done using the *heat-tree* command in GAPPA v.0.6.1 (Czech et al., 2020), with labels that were modified in CorelDraw.

### Post placement analyses

The phylogenetic placement of each query sequence into one of the four colpodean subtaxa or the multiple outgroups was visualized based on the top likelihood weight ratio value obtained from the *jplace* file (i.e., the most likely placement location). To determine which queries were phylogenetically placed into each of the four colpodean subtaxa (Colpodida, Cyrtolophosidida, Bursariomorphida, and Platyophryida) and the outgroup, we used the *extract* command in GAPPA. The command extracts the queries that were placed into user-defined clades of the reference tree. A challenge is that one query sequence can be placed into several different subtaxa. To overcome this, we set a minimum percentage of 99% of likelihood mass (accumulated likelihood weight ratios) that needs to be in a given subtaxon clade for a given query. Otherwise, the query is assigned to a separate “uncertain” category. Furthermore, if a query sequence is placed outside of the predefined subtaxa, the algorithm puts this query sequence into the category “basal branches”. However, none of our query sequences were put into one of the latter categories, indicating that all sequences indeed belong to one of the four colpodean clades or the outgroup with high likelihood.

In a second step, the top seven likelihood weight ratio values in the *jplace* file obtained from phylogenetic placement were summed up for each query sequence using the *assign* command with the *per-query* option in GAPPA. The *assign* command assigns the likelihood weight ratio values of each placement to the user-defined taxonomic groups, and the *per-query* option calculates the sum of these values for each OTU (query). Since likelihood weight ratios were calculated for multiple placements on several different branches, we wanted to show how much support a query sequence had for the respective subtaxon into which it was placed.

To calculate the expected distances between the top seven placement locations of each query sequence, we used the *edpl* command in GAPPA (Matsen et al., 2010). The expected distance between placement locations shows how far the different placement locations of each query are spread out from each other across the branches, weighted by their respective likelihood weight ratios. Higher distances between placement locations of a query might indicate less certainty or accuracy of a sequence being placed in a specific neighbourhood of the reference tree, or an inappropriate reference tree with the incorrect ingroups and outgroups; smaller distances might indicate a higher placement certainty into a small neighbourhood of the tree.

## RESULTS

### OTUs and pairwise comparisons

The re-cleaning and re-clustering of the environmental metabarcoding data of Mahé et al. (2017) with updated programs, yielded 326 OTUs that were assigned to the Colpodea using pairwise alignment comparisons. All of these OTUs were further assigned to one of the four main subtaxa of the Colpodea: 15 to Bursariomorphida, 227 to Colpodida, 54 to Cyrtolophosidida, and 30 to Platyophryida. The maximum pairwise similarity value to the closest reference in the PR^2^ database was 100.0 % for seven OTUs, and the minimum similarity value was 70.5 % for one OTU (**Figure 1a**). The mean similarity value for the OTUs to the reference database was 96.05 %.

**FIGURE 1:**
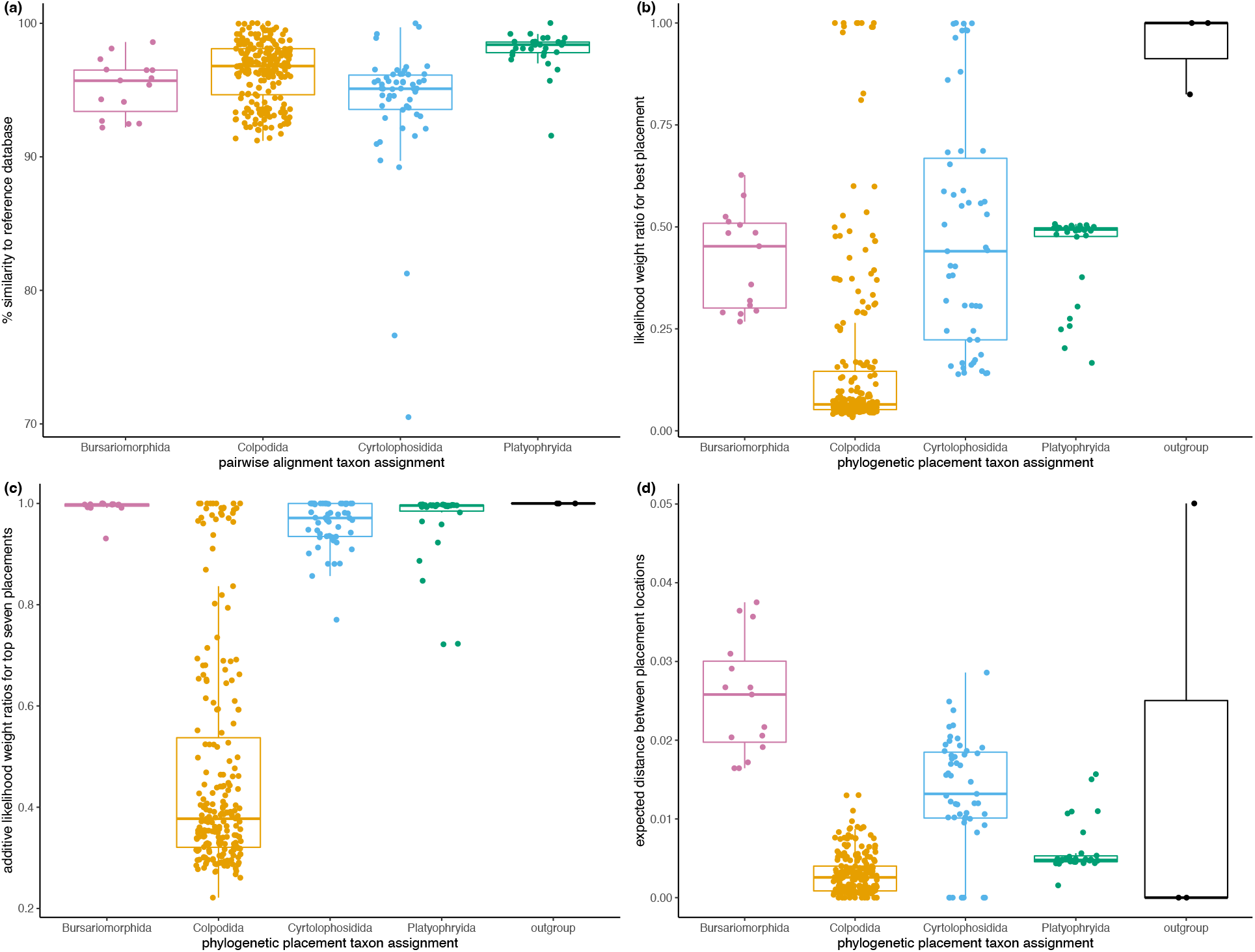
Taxonomic assignment of the colpodean ciliate OTUs to one of the four colpodean subtaxa, Bursariomorphida, Colpodida, Cyrtolophosidida, and Platyophryida, or the outgroup. **(a)** Taxonomic assignment according to the percentage similarity of each OTU to the closest reference in the PR2 database using pairwise alignments. **(b)** Taxonomic assignment according to the top likelihood weight ratio value of each OTU for its best phylogenetic placement on the reference tree. **(c)** Taxonomic assignment according to the sum of the likelihood weight ratio values of each OTU for its top seven phylogenetic placements on the reference tree. **(d)** Expected distance (in branch length units) between the (top seven) phylogenetic placement locations of each OTU in the four different subtaxa and outgroup.

### Evaluating placements on the reference tree

Out of the 326 OTUs that were assigned to the Colpodea by pairwise alignments, 323 OTUs were phylogenetically placed on branches within the Colpodea (**Figure 2**). It should be noted that the heat-trees can visualize more than one potential placement location of an OTU, if the likelihood weight ratio scores (placement probabilities) are distributed across multiple branches. In other words, the heat tree shows the accumulated likelihood weight ratios of all placed sequences on each branch, which can hence be interpreted as an accumulated probability distribution of the placed sequences In our analyses, we used EPA-ng’s default parameter of reporting up to seven most likely placements per sequence, although in theory some OTUs may have non-zero (or close to zero) placement probabilities on more than seven branches (when there are more than seven placements, the likelihood weight ratio scores of these additional placements, however, tend to be very low, at least in our data where the reference tree captures the diversity of the sequences well). The pairwise similarity values for the 323 OTUs that EPAng placed within the Colpodea were 89.2 % or higher (**Figure 1b**). The 323 OTUs that were placed within the Colpodea were also placed within the same subtaxa they were assigned to via pairwise alignments, or on the branch leading up to the clade composed of that subtaxon. The most probable placement location of each OTU always fell into its subclade, although in some cases, some of the other likely placement locations were on branches leading up to their respective subclades. Three OTUs, which were assigned to the Cyrtolophosidida using pairwise alignment comparisons, phylogenetically placed on branches in the outgroup; the pairwise similarity values for these OTUs that placed in the outgroup were also the three lowest values: 70.5 %, 76.6 %, and 81.3 %.

**FIGURE 2:**
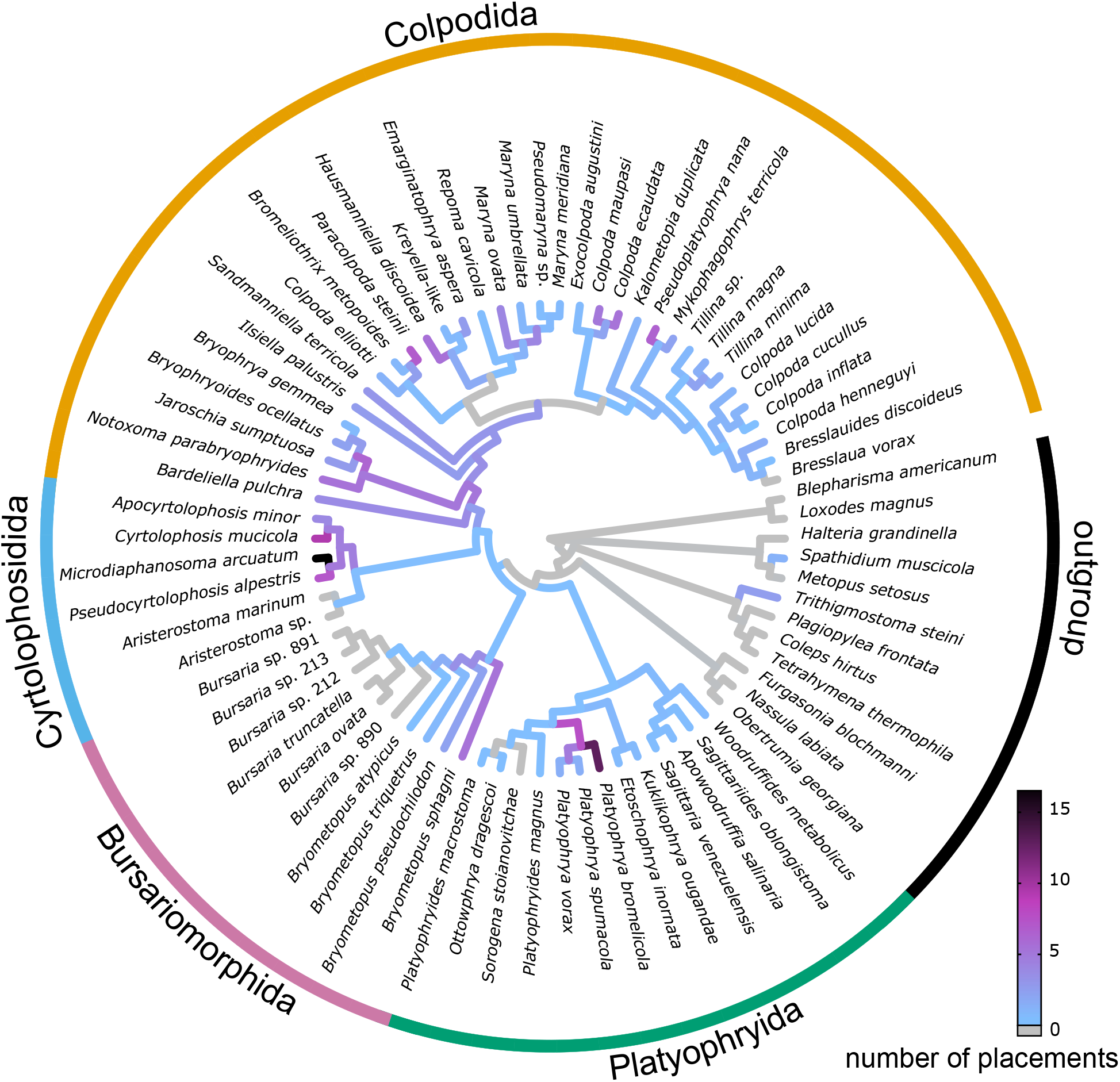
Phylogenetic placements of the colpodean ciliate OTUs across the colpodean reference tree, visualized as a heat tree showing the distribution of placements: no placements occurred on grey branches, whereas coloured branches indicate that sequences were placed on the respective branch with some probability. The “number of placements” per branch is the accumulated sum of likelihood weight ratios of all placements that had non-zero placement probability on that branch.

Of the 326 OTUs, 19 were placed on a single branch with a likelihood weight ratio of almost 1.0. The other 307 OTUs had multiple placements on different branches with varying likelihood weight ratio scores. In the Bursariomorphida, mean likelihood weight ratio score = 0.42, max = 0.63, and min = 0.27. In the Colpodida, mean = 0.16, max = 1.0, and min = 0.03. In the Cyrtolophosidida, mean = 0.48, max = 1.0, and min = 0.14. In the Platyophryida, mean = 0.44, max = 0.51, and min = 0.17. The three OTUs that were placed in the outgroup had a max likelihood weight ratio score of 1.0, 1.0, and 0.83, respectively, indicating that their phylogenetic placement is rather confident.

To obtain more certainty about whether an OTU falls into a specific subtaxon, we summed the top seven likelihood weight ratios of the different placement locations within the subtaxon that each OTU was placed into (**Figure 1c**). This way, by accumulating placement probabilities within subtaxa, the placement uncertainty is taken into account. As in all cases, the taxonomic assignment of all seven reported placement locations of each sequence was the same, accumulating the likelihood weight ratios of these seven locations hence reveals the overall probability of each sequence being placed into their respective clade. In addition to the 19 OTUs that had only one placement on the tree with a likelihood weight ratio score of 1.0, ten more OTUs reached a value of almost 1.0 when their top seven likelihood weight ratios for distinct placements were accumulated. The likelihood weight ratio mean, maximum, and minimum values across all OTUs increased in all subtaxa as follows: In the Bursariomorphida, mean score = 0.62, max = 1.0, and min = 0.93. In the Colpodida, mean score = 0.47, max = 1.0, and min = 0.22. In the Cyrtolophosidida, mean score = 0.96, max = 1.0, and min = 0.77. In the Platyophryida, mean score = 0.96, max = 0.998, and min = 0.72. In the outgroup comprised of several sequences, the one OTU that had more than one likely placement location, reached a likelihood weight ratio value of 1.0 when adding up the values for its placement locations. The OTUs that were placed into the Colpodida still had the lowest likelihood weight ratios when summing over their top seven values. The mean across all OTUs increased by 0.31 when adding up the top seven compared to just the max likelihood weight ratio score, meaning that on average, 31% of the probability distribution of the sequences was allocated to placement locations other than the top most likely one. In the other three subtaxa, the mean increased more when adding up the values for the different top seven placements. In the Bursariomorphida, the mean increased by 0.57, in the Cyrtolophosidida, it increased by 0.48, and in the Platyophryida, the mean increased by 0.52. The Platyophryida, however, was the only subtaxon for which the top seven values did not add up to 1.0 for any OTU, indicating that placements into that clade are less certain as to which specific branch is the most likely placement location.

Since one OTU can have multiple likely phylogenetic placements where the likelihood weight ratio scores are spread across multiple branches, we calculated the expected distance between placement locations (EDPL measured in branch length units, Matsen et al., 2010) for the phylogenetic placements (**Figure 1d**). When distances between the multiple placement locations of an OTU along the branches of the reference tree are low, even if all likelihood weight ratio scores are also low, it increases our confidence of the OTU falling into a specific region of the tree. On the other hand, if the distances between the placement locations are high, our confidence of the assignment of the OTU to a specific subtaxon is decreased, because it indicates that several likely placement locations are spread across the tree.

Of the 326 OTUs from this dataset, the expected distance between placement locations has a mean of 0.01, which is relatively low compared to the average branch length of the tree. This indicates that, on average, the placement distribution of an OTU is spread across neighbouring branches of the tree. For 262 of these OTUs, the EDPL was < 0.01, and for 63 OTUs, it ranged between 0.01 and 0.038, while one OTU, which was placed on two branches in the outgroup, had the highest value of 0.05. In the Bursariomorphida, for the expected distance between placement locations we obtained mean = 0.025, max = 0.04, min = 0.02. In the Colpodida, where the highest likelihood weight ratio scores of the OTUs were lowest among all subtaxa we obtained mean = 0.0, max = 0.1, and min = 0.0. In the Cyrtolophosidida, the mean = 0.01, max = 0.03, and min = 0.0. In the Platyophryida, we obtained mean = 0.0, max = 0.02 and min = 0.0. These low expected distance values of the between placements show that even if there are multiple placements with low likelihood weight ratio scores, the placements are very close to each other in the reference tree.

### Placements on specific taxonomic branches

The OTUs were largely placed throughout the colpodean reference tree, although there were no placements on some of the branches (Figure 2). The branches with the most placements were those that led to *Microdiaphanosoma arcuatum* and *Platyophrya bromelicola*. Multiple OTUs were placed on the branches leading up to *Mykophagophrys terricola* and *Pseudoplatyophrya nana*. No OTUs were placed on the branch leading to *Bursaria truncatella* or *Sorogena stoianovitchae*.

## DISCUSSION

Using environmental metabarcoding data from colpodean ciliate OTUs obtained from Neotropical rainforest soils, we found three main results. First, there can be differences in taxonomic assignments of the OTUs between pairwise alignments and phylogenetic placements, when the pairwise similarities of the OTUs to the reference database are low. Second, when there were multiple likely placement locations for an OTU, the summed up likelihood weights of the top seven placement locations, as well as the distance between those locations, can provide additional support for the interpretation of the placement of the OTU into a subtaxon. Third, the colpodean ciliate OTUs from the Neotropical soils were distributed over the entire reference tree, with most placements on branches leading to species found in soils and ground waters around the world.

The taxonomic assignments in this study were inferred by pairwise comparisons as implemented in VSEARCH, and by phylogenetic placements as implemented in EPA-ng. There are other programs that conduct these, and other types, of taxonomic assignments (Hleap et al., 2021). We employed VSEARCH and EPA-ng because they are both widely used in analysing environmental sequencing datasets of protists, such as ciliates, and because of the extensive power of the post-processing options for phylogenetic placement results implemented in GAPPA. We chose not to optimize any of the programs’ parameters, because deriving them from mock communities (e.g., Hleap et al., 2021) is not yet feasible for most natural protistan communities, such as those from Neotropical rainforest soils. The level of taxonomic assignments that we focused on here was above the species level. Species-level taxonomic assignments for protistan environmental sequencing datasets are not yet feasible because of the lack of sufficient identified species in the reference database in general (Berney et al., 2017; Guillou et al., 2013), and in colpodean species in specific (Rajter et al., 2021).

### Differences and similarities between pairwise alignments and phylogenetic placements

Out of the 326 OTUs assigned to the Colpodea using pairwise alignments, three OTUs were phylogenetically placed in the outgroup; these three OTUs had the lowest pairwise similarity to the PR2 database (70.5%, 76.6%, and 81.3%, respectively). We could not taxonomically identify these three OTUs with any confidence based on the reference tree we used, since there was only a limited selection of other major ciliate taxa in the outgroup. Jamy et al. (2020) did also observe a disagreement between their pairwise alignment and comprehensive phylogenetically-informed analyses, but here we showed that the disagreement can also arise just between pairwise alignments and phylogenetic placements.

For the three OTUs where there was disagreement on the taxonomic assignment between pairwise alignments and phylogenetic placements, we did not evaluate the cause of the disagreement nor can we say which approach is more accurate, as much more analyses are needed to do that in depth. However, the differences underlying either the pairwise alignment or the phylogenetic placement approaches may be related to the debate around the underlying approaches to distance-versus character-based methods for inferring phylogenies (Felsenstein, 2004). With the pairwise alignment approach, we compared an overall similarity without taking into account evolutionary processes, such that we could have been misled by not using an explicit evolutionary model. By contrast, the phylogenetic placement approach takes evolutionary processes into account, which however also means that we could have been misled by using such a (relatively) parameter-rich model. The phylogenetic placement approach could have also been misled because it was placing short metabarcoding reads with too little phylogenetic signal, but at least in ciliates, the V4 fragment does have a relatively strong phylogenetic signal compared to other short fragments of the SSU-rRNA locus (Berger et al., 2011; Dunthorn et al., 2014).

The remaining 323 OTUs were assigned to the same colpodean subtaxa by both pairwise alignments and phylogenetic placements. Since the taxonomic assignment of these 323 OTUs to the four different subtaxa was the same with both pairwise alignments and phylogenetic placements, high pairwise similarity values indicate a higher likelihood of no differences between the methods. The taxonomic assignments were only evaluated at the level of the four main subtaxa of the Colpodea: i.e., either to the Bursariomorphida, Colpodida, Cyrtolophosidida, or Platyophryida. These four subtaxa all have support for their monophyly from nuclear and mitochondrial loci (Dunthorn et al., 2008, 2011; Foissner et al., 2011; Rajter et al., 2021; Vd’ačný & Foissner, 2019). The reason why the taxonomic assignments were not evaluated at higher taxonomic resolution is that within these subtaxa, there is little phylogenetic resolution as many of the known smaller clades and most known species have yet to be sequenced (Dunthorn et al., 2008, 2012; Foissner et al., 2014; Rajter et al., 2021). Given these insights about colpodean ciliates, it is useful to know about the phylogenetic support for the taxa that one intends to identify using phylogenetic placement methods. In other words, when using phylogenetic placements for taxonomic assignment, this is not just a bioinformatics problem, but also a problem that requires taxonomic knowledge.

### Additive likelihood values and distances between placements

Many of the colpodean OTUs analysed here had multiple likely placement locations on the reference tree. These alternative placements were not considered in most published phylogenetic placement studies using EPA-ng, (Aguirre-von-Wobeser, 2021; Bass et al., 2018; Fuchsman et al., 2022; Gottschling et al., 2021; Iniesto et al., 2021; Metz et al., 2021; Schoenle et al., 2020). Here, for the first time, we showcased together the features of the program GAPPA tool that can take multiple likely placements when performing taxonomic assignments into account.

One way to analyse OTUs when they have multiple placements is to use GAPPA to sum up the likelihood weight ratio values of the top *n* (here, *n* = 7) placement locations within a subtaxon, using the *assign* command with the *per-query* option. More placement locations could have been stored in the resulting *jplace* placement file, but we used the default output of EPA-ng that only reports the top seven. In the Bursariomorphida, Cyrtolophosidida, and Platyophryida, the sums over the likelihood weight ratios were high when summing up the top seven values calculated for the different placements within the subtaxon. This means that the assignment to the subtaxon is supported. In the Colpodida, however, these sums over the likelihood weight ratio values of the top seven placement values were still low after adding them up. This is due to there being more than seven almost equally likely placement locations of an OTU across the branches of this subtaxon. These multiple likely placements may be due to there being many closely related references for the Colpodida in the tree; in future analyses, one could further quantify this by computing the mean and variance of branch lengths in that subtree of the reference compared to the rest of the tree or by conducting a leave-one-out test of placement sequences from that reference clade into the tree again.

Another way to analyse OTUs when they have multiple placements is to use the expected distance between placement locations, as implemented for example in the GAPPA *edpl* command (Czech et al., 2020; Matsen et al., 2010) to estimate the average distances between the different placement locations of an OTU; in future analyses, one could normalize this by comparing it to random placements and then quantifying to which extent they are different from a random placement. The average distances between the different placement locations provide a measure of the spread of the placements across the reference tree; i.e., if there is a local uncertainty between close branches, the distances will be low; while if there is a global uncertainty between dispersed branches, the distances will be high. Again, we only considered the distances between the top seven placements because, but we could have looked at more if we were not using the default parameter that will be implemented by most end-users. The distances between the different placement locations of an OTU were low in all four subtaxa. In the Colpodida, the distances were the lowest among all subtaxa due to there being many closely related reference sequences. This means that the placements within the subtaxon are very close to each other and that the taxonomic assignments to that subtaxon were supported.

Evaluating both the sum of the likelihood weight ratio values of the top seven (or some other number of choice) placements of an OTU and the distances between the different placement locations, are features that can be more widely used in phylogenetic placement analyses of environmental sequencing data, beyond just looking at the most likely placement. Combined, these analyses increase the confidence in the taxonomic assignment of an OTU to a taxon.

### Evolutionary and ecological interpretations of the phylogenetic placements

Following recommendations by Rajter et al. (2021), we can use these phylogenetic placements to tentatively interpret the colpodean OTUs from Neotropical rainforest soils. From an evolutionary perspective, no new subtaxa were found. Although some OTUs were placed on branches leading up to one of the four colpodean subtaxa, these OTUs had multiple placement locations and could be found on other branches within their respective subtaxon with higher likelihood weight scores. We therefore already know the major colpodean groups, at least for those that reside in Neotropical soils. Metabarcoding of soils elsewhere has also not uncovered new colpodean subtaxa (Venter et al., 2018), but marine environments may contain novel groups (Dunthorn et al., 2014; Gimmler et al., 2016).

From an ecological perspective, given that all four major subclades largely contain resting/dormancy cysts, have r-selected reproductive strategies, and are bacterivorous via general filtration of water, we can infer that most of the OTUs that were phylogenetically placed within the Colpodea likewise derive from species with these same characteristics. For example, *M. arcuatum* is a typical ciliate with these characteristics, and has been found worldwide in soils and in groundwaters (Foissner, 1993; Quintela-Alonso et al., 2011). Other OTUs can change their place in a food web, such as those that led to the branch with *P. bromelicola*, which can form small cells that are bacteriovorous and large cells that are predatory on other protists (Foissner & Wolf, 2009). Other OTUs placed on the branches to *M. terricola* and *P. nana*, which are fungivorous with modified oral structures that allow them to feed on hyphae or yeast cells and are found in soils and freshwaters (Foissner, 1993). No OTUs were detected on the branch leading up to *B. truncatella*, which is a predator on other ciliates and has been observed worldwide in freshwaters (Foissner, 1993), nor on the branch leading to *S. stoianovitchae*, which has a multicellular sorocarpic-like life cycle stage and has been isolated in other tropical and temperate environments (Foissner, 1993).

## CONCLUSION

We found that there can be differences in taxonomic assignments of OTUs between pairwise sequence comparison approaches and phylogenetic placement approaches; as expected, these differences are in assignments with a low percent similarity to the reference database, at least in colpodean ciliates. Phylogenetic placements may be better suited for taxonomic assignment because the method incorporates more evolutionary information. We also showed how evaluating the likelihoods and distances when there are multiple likely OTU placements can allow researchers to further evaluate the uncertainty in these taxonomic assignments. With high throughput sequencing, and using long-read sequencing such as PacBio, inferences relying on phylogenetic placement methods are likely to improve in the future because of the increase in molecular data per sequence. The approach developed here to taxonomically identify colpodean ciliate OTUs can be easily applied to other environmental DNA sequencing datasets of both microbes and macrobes.

## ACKNOWLEDGEMENTS

We thank Pierre Barbera and Florian Leese for helpful comments and suggestions. Funding came from the Alexander von Humboldt Foundation to LR.

---

**FIGURE S1.**
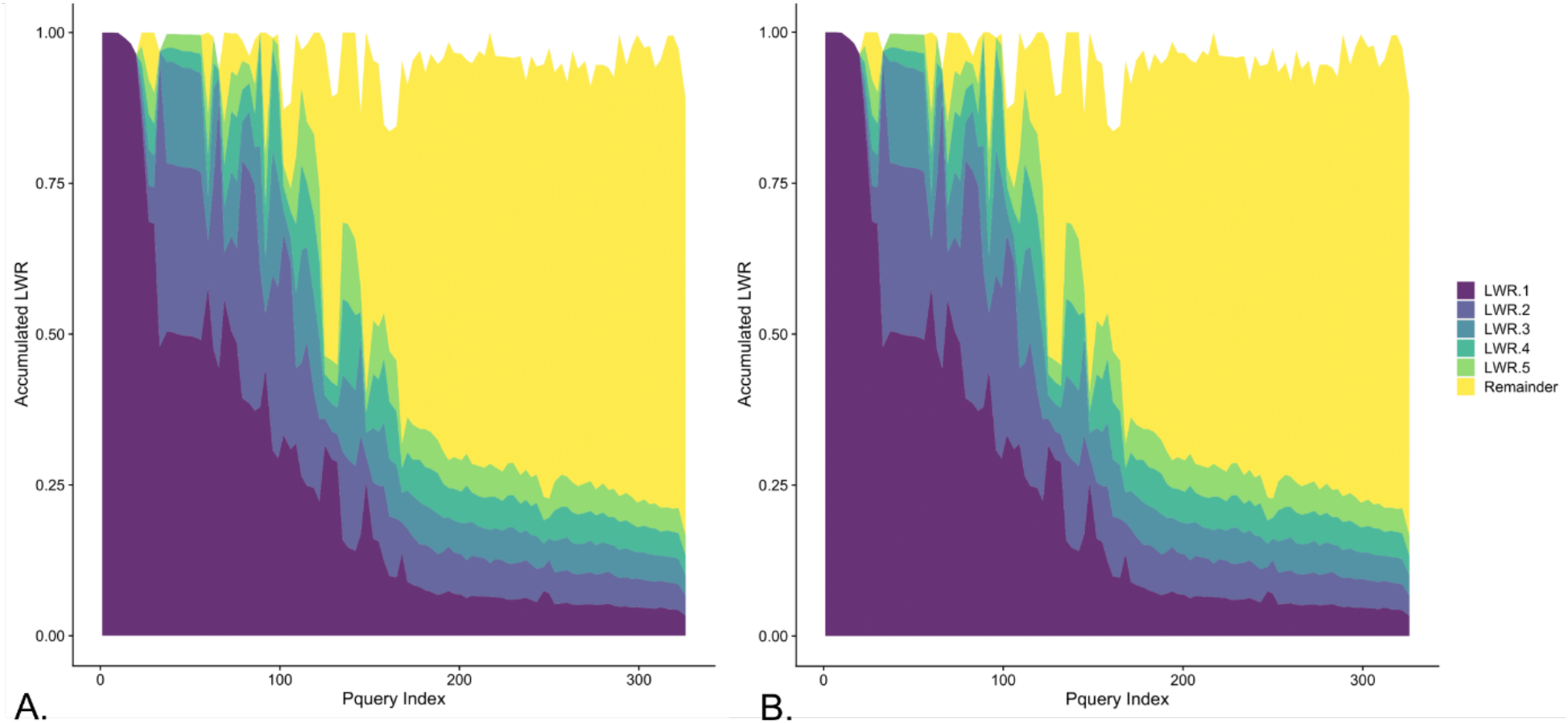
The distribution of the likelihood weight ratios of the top 70 **(A)** and 700 **(B)** placements. On the x-axis are displayed individual query sequences used in the phylogenetic placement. They are sorted in a way that puts more confident placements towards the left hand side. On the y-axis is the accumulated likelihood weight ratio of the placed query sequences. As a single query sequence may have multiple placement locations, the color code differentiates the placements with the highest likelihood weight ratio (LWR.1), the second highest likelihood weight ratio (LWR.2), etc. Only the top five likelihood weight ratios are shown in color; the rest is accumulated in the remainder (yellow). The individual query sequences (on the x-axis) are sorted by decreasing confidence, weighing their top n probabilities with a factor of 1/n each: query sequences with the highest top likelihood weight ratio on the left side to the lowest top likelihood weight ratio on the right side.

## Notes

### Competing Interest Statement

The authors have declared no competing interest.

