## Supplementary File 1 for "Comparison of phylogenetic placements to pairwise alignments for taxonomic assignment of ciliate OTUs": File_S1.html

xml version="1.0" encoding="utf-8"?


Ciliate SSU-rRNA reference alignments and trees for phylogenetic placements

### Table of Contents

- 1. Overview
- 2. amplicon filtering and annotation
  - 2.1. helpers (python scripts)
    - 2.1.1. OTU cleaver
    - 2.1.2. merge sub superstring OTUs with larger OTUs
    - 2.1.3. stampa merge
    - 2.1.4. OTU table updater
- 3. phylogenetic placement

- 4. R scripts

Isabelle Ewers, Ľubomír Rajter, Lucas Czech, Frédéric Mahé, Micah Dunthorn

supplementary file for the bioinformatics methods

### 1 Overview

DNA extraction and high-throughput amplicon Illumina MiSeq sequencing
protocols correspond to Mahé et al. (2017).

| primer name | 5'-3' sequence | length | position | reference |
| --- | --- | --- | --- | --- |
| TAReuk454FWD1 | CCAGCASCYGCGGTAATTCC | 20 | 564 | Stoeck et al (2010) |
| TAReukREV3 | ACTTTCGTTCTTGATYRA | 18 | 963 | Stoeck et al (2010) |

**position**: position of the first nucleotide of the primer in the
reference sequence *Saccharomyces cerevisiae* (FU970071).

Amplicons were processed using the following swarm-based pipeline:
paired-end reads were merged with vsearch's `--fastq_mergepairs`
command (version 2.15.1, allowing for staggered reads), and trimmed
with cutadapt (version 3.0), keeping only reads containing both
forward and reverse primers. After trimming, the expected error per
read was estimated with vsearch's command `--fastq_filter` and the
option `--eeout`. Each sample was then dereplicated, i.e. strictly
identical reads were merged, using vsearch's command
`--derep_fulllength`, and converted into fasta format. Clustering was
performed at the sample level with swarm using default parameters, and
results were set aside for later use.

To prepare for a global clustering, individual fasta files (one per
sample) were pooled and further dereplicated with vsearch. Files
containing per-read expected error values were also dereplicated to
retain only the lowest expected error for each unique sequence. Global
clustering was performed with swarm (using the `--fastidious`
option). Cluster representative sequences were then searched for
chimeras with vsearch's command `--uchime_denovo` (Edgar et al.,
2011). In parallel, representative sequences were assigned to taxa
using vsearch's `--usearch_global` command, and EukRibo v2020-10-27, a
customized version of the reference database PR2.

Clustering results, expected error values, taxonomic assignments, and
chimera detection results were used to build a "raw" occurrence table,
where only reads without primers, reads shorter than 32 nucleotides
and reads with uncalled bases ("N") were discarded. In order to build
a "filtered" occurrence table, additional filters were applied on
clusters and cluster representative sequences. Non-chimeric sequences,
sequences with an expected error per nucleotide below 0.0002, and
clusters containing at least 2 reads were retained. Primer trimming is
not perfect and some sequences can still contain primer fragments or
be excessively trimmed. These sub- or super-sequences were identified
using vsearch and merged with their closest, most abundant perfectly
trimmed sequence. Finally, occurrence patterns throughout our sample
collection were used to refine our results. Some clusters contain
sub-clusters with only a single-nucleotide difference but with
different ecological patterns, here defined as uncorrelated abundance
values in at least 5% of the samples. These sub-clusters were turned
into distinct clusters (see the 'OTU cleaving' python script
below). On the other hand, some clusters with similar sequences
present correlated abundance values in at least 95% of the
samples. These correlated clusters were merged using mumu, a C++
re-implementation of lulu's method (Frøslev et al. 2017).

A reference alignment and a tree were produced for the ciliates. All
environmental sequences with a similarity higher ciliates than any
other clade were extracted from the Neotropical dataset and placed on
the ciliate multiple sequence alignment using EPA-ng, PaPaRa, and
gappa.

### 2 amplicon filtering and annotation

Raw fastq files available in NCBI/EBI bioproject
PRJNA317860. Occurrence table and fasta files available on demand.

```
#!/bin/bash
cd ${HOME}/projects/Neotropical_Soils/data/

## ------------------------------------------------- adapt variables to project

ENCODING="33"
THREADS=4
PRIMER_F="CCAGCASCYGCGGTAATTCC"
PRIMER_R="ACTTTCGTTCTTGATYRA"
ANTI_PRIMER_R="TYRATCAAGAACGAAAGT"
VSEARCH="$(which vsearch) --quiet"
CUTADAPT="$(which cutadapt) --minimum-length 32 --cores=${THREADS} --discard-untrimmed --times=2"
SWARM="$(which swarm)"
RESOLUTION=1

## ------------------------------------------------------------- fastq to fasta
echo "from fastq to fasta files..."
export LC_ALL=C

MIN_F=$(( ${#PRIMER_F} * 2 / 3 ))  # primer match is >= 2/3 of primer length
MIN_R=$(( ${#PRIMER_R} * 2 / 3 ))
FIFOS=$(echo fifo_{merged_fastq,trimmed_fastq,filtered_fastq})

# summarize read quality (error rates)
for PAIR in 1 2 ; do
    zcat ./*_[1-9]_${PAIR}.fastq.gz | \
        ${VSEARCH} \
            --fastq_eestats - \
            --output R${PAIR}_eestats.log &
done

i=0
for FORWARD in *_[1-9]_1.fastq.gz ; do
    i=$(( $i + 1 ))
    echo -e "$i\t${FORWARD}"
    REVERSE="$(sed -r 's/_([1-9])_1/_\1_2/' <<< "${FORWARD}")"
    SAMPLE="${FORWARD/_lib*.fastq.gz/}"
    LOG="${SAMPLE}.log"
    FASTA="${SAMPLE}.fas"
    QUAL="${SAMPLE}.qual"
    LOCAL_STATS="${SAMPLE}.stats"

    # clean
    rm -f ${SAMPLE}.{fas,qual,log,stats} ${FIFOS}
    mkfifo ${FIFOS}

    # merge
    (${VSEARCH} \
         --threads "${THREADS}" \
         --fastq_mergepairs "${FORWARD}" \
         --reverse "${REVERSE}" \
         --fastq_ascii "${ENCODING}" \
         --fastq_allowmergestagger \
         --fastqout fifo_merged_fastq 2> "${SAMPLE}.log" &
    )

    # trim
    (${CUTADAPT} \
         --revcomp \
         -g "${PRIMER_F}" \
         -O "${MIN_F}" fifo_merged_fastq 2>> "${LOG}" | \
         ${CUTADAPT} \
             -a "${ANTI_PRIMER_R}" \
             -O "${MIN_R}" - > fifo_trimmed_fastq 2>> "${LOG}" &
    )

    # convert fastq to fasta, and extract error rate
    (${VSEARCH} \
         --fastq_filter fifo_trimmed_fastq \
         --fastq_maxns 0 \
         --relabel_sha1 \
         --fastq_ascii "${ENCODING}" \
         --eeout \
         --fasta_width 0 \
         --fastaout - 2>> "${LOG}" | \
         tee >(paste - - | awk -F "[>;=\t]" '{print $2, $4, length($NF)}' | \
                   sort -k3,3n -k1,1d -k2,2n | \
                   uniq --check-chars=40 > "${QUAL}") > fifo_filtered_fastq &
    )

    # dereplicate
    ${VSEARCH} \
        --derep_fulllength fifo_filtered_fastq \
        --sizeout \
        --fasta_width 0 \
        --xee \
        --output "${FASTA}" 2>> "${LOG}"

    # list local clusters
    ${SWARM} \
        --threads "${THREADS}" \
        --differences "${RESOLUTION}" \
        --usearch-abundance \
        --log /dev/null \
        --output-file /dev/null \
        --statistics-file - \
        "${FASTA}" | \
        awk 'BEGIN {FS = OFS = "\t"} $2 > 2' > "${LOCAL_STATS}"

    # clean
    rm ${FIFOS}
done


## ---------------------------------------------------------- global clustering
echo "run global clustering and chimera detection..."
cd ${HOME}/projects/Neotropical_Soils/results/

## variables and lists of files
FASTA_FILES=$(find ../data/2013-12-30_reduced/ -name "*.fas" -type f ! -empty | \
                  tr "\n" " " | sed 's/\n$//')
QUALITY_FILES=$(sed 's/.fas/.qual/g' <<< ${FASTA_FILES})
STATS_FILES=$(sed 's/.fas/.stats/g' <<< ${FASTA_FILES})
N_SAMPLES=$(tr " " "\n" <<< ${FASTA_FILES} | wc -l)
PROJECT="Neotropical_forest_soils"
FINAL_FASTA="${PROJECT}_18S_V4_${N_SAMPLES}_samples.fas"
QUALITY_FILE="${FINAL_FASTA/.fas/.qual}"
DISTRIBUTION_FILE="${FINAL_FASTA/.fas/.distr}"
POTENTIAL_SUB_SEEDS="${FINAL_FASTA/.fas/_per_sample_OTUs.stats}"
FILTER=2  # exclude clusters with an abundance of 1


## Build expected error file
echo "Pooling expected error observations..."
sort -k3,3n -k1,1d -k2,2n --merge ${QUALITY_FILES} | \
    uniq --check-chars=40 > "${QUALITY_FILE}" &


## Build distribution file (sequence <-> sample relations)
echo "Listing sequence to sample relations..."
for f in ${FASTA_FILES} ; do
    grep -H "^>" "${f}"
done | \
    sed 's/.*\/// ; s/.fas:>/\t/ ; s/;size=/\t/ ; s/;$//' | \
    awk 'BEGIN {FS = OFS = "\t"} {print $2, $1,$3}' > "${DISTRIBUTION_FILE}" &


## list local cluster seeds of size > 2 (some files can be empty)
echo "list local cluster seeds of size > 2..."
for LOCAL_STATS in ${STATS_FILES} ; do
    NAME="${LOCAL_STATS/*\//}"
    NAME="${NAME/.stats/}"
    awk -v NAME="${NAME}" \
        'BEGIN {FS = OFS = "\t"} {print NAME, $0}' "${LOCAL_STATS}"
    unset NAME
done > "${POTENTIAL_SUB_SEEDS}" &


## global dereplication
${VSEARCH} \
    --derep_fulllength <(cat ${FASTA_FILES}) \
    --sizein \
    --sizeout \
    --fasta_width 0 \
    --output "${FINAL_FASTA}"


## clustering
OUTPUT_SWARMS="${FINAL_FASTA%.*}_${RESOLUTION}f.swarms"
OUTPUT_LOG="${FINAL_FASTA%.*}_${RESOLUTION}f.log"
OUTPUT_STATS="${FINAL_FASTA%.*}_${RESOLUTION}f.stats"
OUTPUT_STRUCT="${FINAL_FASTA%.*}_${RESOLUTION}f.struct"
OUTPUT_REPRESENTATIVES="${FINAL_FASTA%.*}_${RESOLUTION}f_representatives.fas"
${SWARM} \
    --differences "${RESOLUTION}" \
    --fastidious \
    --usearch-abundance \
    --threads "${THREADS}" \
    --internal-structure "${OUTPUT_STRUCT}" \
    --output-file "${OUTPUT_SWARMS}" \
    --statistics-file "${OUTPUT_STATS}" \
    --seeds - \
    "${FINAL_FASTA}" 2>> "${OUTPUT_LOG}" | \
    ${VSEARCH} \
        --sortbysize - \
        --fasta_width 0 \
        --output "${OUTPUT_REPRESENTATIVES}"


## fake taxonomic assignment
grep "^>" ${FINAL_FASTA/.fas/_1f_representatives.fas} | \
    sed -r 's/^>//
            s/;size=/\t/
            s/;?$/\t0.0\tNA\tNA/' > ${FINAL_FASTA/.fas/_1f_representatives.results}


## chimera detection
## discard sequences with an abundance lower than FILTER
## filter out low abundant amplicons (if need be) and search for
## chimeras
FILTER=2
${VSEARCH} \
    --fastx_filter "${OUTPUT_REPRESENTATIVES}"  \
    --minsize "${FILTER}" \
    --fastaout - | \
    ${VSEARCH} \
        --uchime_denovo - \
        --log "${OUTPUT_REPRESENTATIVES%%.*}.log" \
        --uchimeout "${OUTPUT_REPRESENTATIVES%%.*}.uchime"


## ------------------------------------------------------------------- cleaving
echo "run cleaving..."
PROJECT="${FINAL_FASTA/.fas/}"
SRC="${HOME}/src"
OTU_CLEAVER="OTU_cleaver.py"

# split OTUs
python3 "${SRC}/${OTU_CLEAVER}" \
        --global_stats "${PROJECT}_1f.stats" \
        --per_sample_stats "${PROJECT}_per_sample_OTUs.stats" \
        --struct "${PROJECT}_1f.struct" \
        --swarms "${PROJECT}_1f.swarms" \
        --fasta "${PROJECT}.fas"

# chimera detection and taxonomic assignment
QUERY1="${PROJECT}_1f_representatives.fas"
QUERY2="${QUERY1}2"
OUTPUT="${QUERY1/.fas/.uchime2}"
LOWEST_ABUNDANCE=$(sed -rn '/^>/ s/.*;size=([0-9]+);?/\1/p' ${QUERY2} | sort -n | head -n 1)

## fake taxonomic assignment
grep "^>" ${QUERY2} | \
    sed -r 's/^>//
            s/;size=/\t/
            s/;?$/\t0.0\tNA\tNA/' > ${QUERY2/.fas2/.results2}

# sort and filter by abundance (default to an abundance of 1), search
# for chimeras
cat ${QUERY1} ${QUERY2} | \
    ${VSEARCH} \
        --sortbysize - \
        --sizein \
        --minsize ${LOWEST_ABUNDANCE:-1} \
        --sizeout \
        --output - | \
    ${VSEARCH} \
        --uchime_denovo - \
        --uchimeout "${OUTPUT}"


## ------------------------------------------------------------ first OTU table
echo "build first OTU table..."
SCRIPT="${SRC}/OTU_contingency_table_filtered.py"
FASTA="${FINAL_FASTA}"
REPRESENTATIVES="${FASTA/.fas/_1f_representatives.fas}"
STATS="${FASTA/.fas/_1f.stats}"
SWARMS="${FASTA/.fas/_1f.swarms}"
UCHIME="${FASTA/.fas/_1f_representatives.uchime}"
ASSIGNMENTS="${FASTA/.fas/_1f_representatives.results}"
QUALITY="${FASTA/.fas/.qual}"
DISTRIBUTION="${FASTA/.fas/.distr}"
OTU_TABLE="${FASTA/.fas/.OTU.filtered.cleaved.table}"

# build OTU table
python3 \
    "${SCRIPT}" \
    --representatives <(cat "${REPRESENTATIVES}"{,2}) \
    --stats <(cat "${STATS}"{,2}) \
    --swarms <(cat "${SWARMS}"{,2}) \
    --chimera <(cat "${UCHIME}"{,2}) \
    --quality "${QUALITY}" \
    --assignments <(cat "${ASSIGNMENTS}"{2,}) \
    --distribution "${DISTRIBUTION}" > "${OTU_TABLE}"


## ------------------------------------------------- merge sub- or superstrings
echo "merge sub- and super-strings..."
PYTHON_SCRIPT="merge_sub_superstring_OTUs_with_larger_OTUs.py"
OUTPUT_TABLE="${OTU_TABLE/.table/.nosubstringOTUs.table}"
TMP_TABLE=$(mktemp)
TMP_UC=$(mktemp)

# extract fasta and search for identical sequences (terminal gaps excepted)
awk 'NR > 1 {printf ">"$1"\n"$10"\n"}' "${OTU_TABLE}" | \
    ${VSEARCH} \
        --threads ${THREADS} \
        --cluster_smallmem - \
        --id 1.0 \
        --qmask none \
        --usersort \
        --uc - | \
    grep "^H" > "${TMP_UC}"

# merge OTUs that are sub- or superstrings of more abundant OTUs
python3 "${SRC}/${PYTHON_SCRIPT}" -t "${OTU_TABLE}" -m "${TMP_UC}" -o "${TMP_TABLE}"

# sort OTUs (using the pre-merging ranking)
(head -n 1 "${TMP_TABLE}"
 tail -n +2 "${TMP_TABLE}" | sort -k1,1n) > "${OUTPUT_TABLE}"

# check number of reads that stay the same
awk 'NR > 1 {total += $2} END {print total}' "${OTU_TABLE}"
awk 'NR > 1 {total += $2} END {print total}' "${OUTPUT_TABLE}"
# count deleted OTUs
wc -l "${OUTPUT_TABLE}" "${OTU_TABLE}"

# clean
rm "${TMP_TABLE}" "${TMP_UC}"


## ------------------------------------------------------------- mumu (ex-lulu)
echo "run mumu..."
MUMU="${HOME}/src/mumu/mumu"
MATCH_LIST="${OUTPUT_TABLE/.table/.match_list}"
REDUCED_TABLE="${OUTPUT_TABLE/.table/_reduced.table}"
LOG="${OUTPUT_TABLE/.table/.mumu.log}"
NEW_TABLE="${OUTPUT_TABLE/.table/_lulu.table}"

# extract fasta sequences from OTU table
awk 'NR > 1 {printf ">"$4";size="$2";\n"$10"\n"}' "${OUTPUT_TABLE}" \
   > "${OUTPUT_TABLE/.table/.fas}"
cut -f 4,14- "${OUTPUT_TABLE}" > "${REDUCED_TABLE}"

# find similar sequences, discard abundance values
${VSEARCH} \
    --usearch_global "${OUTPUT_TABLE/.table/.fas}" \
    --db "${OUTPUT_TABLE/.table/.fas}" \
    --self  \
    --id 0.84 \
    --iddef 1 \
    --userfields query+target+id \
    --maxaccepts 0 \
    --query_cov 0.9 \
    --maxhits 10 \
    --userout - | \
    sed -r 's/;size=[0-9]+;//g' > "${MATCH_LIST}"

## run mumu
${MUMU} \
    --otu_table "${REDUCED_TABLE}" \
    --match_list "${MATCH_LIST}" \
    --new_otu_table "${NEW_TABLE}" \
    --log "${LOG}"


## Taxonomic assignment, search for best hits (eliminate chimeras and singletons first)
echo "taxonomic assignment..."
QUERY="${OUTPUT_TABLE/.table/.fas}"
RESULTS="${QUERY/.fas/.results}"
DATABASE="${HOME}/data/references/46345_EukRibo_V4_2020-10-22.fas"

${VSEARCH} \
    --usearch_global "${QUERY}" \
    --threads ${THREADS} \
    --db "${DATABASE}" \
    --dbmask none \
    --qmask none \
    --rowlen 0 \
    --notrunclabels \
    --userfields query+id1+target \
    --maxaccepts 0 \
    --maxrejects 32 \
    --top_hits_only \
    --output_no_hits \
    --id 0.5 \
    --iddef 1 \
    --userout - | \
    sed 's/;size=/_/ ; s/;//' > hits.representatives

## in case of multi-best hit, find the last-common ancestor
python3 ${HOME}/src/stampa_merge.py $(pwd)

## sort by decreasing abundance
sort -k2,2nr -k1,1d results.representatives > "${RESULTS}"

## clean
rm hits.representatives results.representatives


## ------------------------------------------------------ build final OTU table
echo "build final OTU table..."
PYTHON_SCRIPT="OTU_table_updater.py"
PATH_TO="$(pwd)"
OLD_TABLE="${OUTPUT_TABLE/.table/_lulu.table}"
NEW_TAXONOMY="${OLD_TABLE/.table/.results}"
NEW_TABLE=$(mktemp)

python3 \
    "${SRC}/${PYTHON_SCRIPT}" \
    --old_otu_table "${PATH_TO}/${OLD_TABLE}" \
    --new_taxonomy "${PATH_TO}/${NEW_TAXONOMY}" \
    --new_otu_table "${NEW_TABLE}"

mv "${NEW_TABLE}" "${PATH_TO}/${OLD_TABLE}" && chmod go+r,g+w "${PATH_TO}/${OLD_TABLE}"


## --------------------- extract representative sequences assigned to ciliates

FASTA="${NEW_TABLE/.table/.Ciliophora.fas}"
grep "Ciliophora" "${NEW_TABLE}" | \
    awk '{printf ">"$4";size="$2"; "$11" "$12"\n"$10"\n"}' > "${FASTA}"

FASTA="${TABLE/.table/.Colpodea.fas}"
grep "Colpodea" "${NEW_TABLE}" | \
    awk '{printf ">"$4";size="$2"; "$11" "$12"\n"$10"\n"}' > "${FASTA}"

exit 0
```

#### 2.1 helpers (python scripts)

Helper python scripts listed below are also available at
https://github.com/frederic-mahe/fred-metabarcoding-pipeline, in the
`src` folder.

##### 2.1.1 OTU cleaver

```
#!/usr/bin/env python3
# -*- coding: utf-8 -*-
"""
   break swarm clusters, using sample distribution data
"""

__author__ = "Frédéric Mahé <>"
__date__ = "2021/03/19"
__version__ = "$Revision: 1.2"

import os
import re
import sys
import copy
import argparse
import operator


# *************************************************************************** #
#                                                                             #
#                                  Functions                                  #
#                                                                             #
# *************************************************************************** #

if __name__ == '__main__':
    """
    Parse arguments from command line.
    """
    parser = argparse.ArgumentParser(
        description="break swarm clusters, using sample distribution data.")

    parser.add_argument("--global_stats",
                        dest="global_stats_file",
                        required=True,
                        help="cluster statistics")

    parser.add_argument("--per_sample_stats",
                        dest="per_sample_stats_file",
                        required=True,
                        help="per-sample cluster statistics")

    parser.add_argument("--fasta",
                        dest="fasta_file",
                        required=True,
                        help="amplicon sequences")

    parser.add_argument("--struct",
                        dest="struct_file",
                        required=True,
                        help="internal structure of clusters")

    parser.add_argument("--swarms",
                        dest="swarms_file",
                        required=True,
                        help="list of amplicons per cluster")

    ARGS = parser.parse_args()


def per_sample_stats_parse(per_sample_stats_file, percentage):
    """
    Map samples, OTU seeds and stats.
    """
    separator = "\t"
    per_sample_stats = [dict() for i in range(0, 256)]
    number_of_samples = 0
    previous_sample = None

    with open(per_sample_stats_file, "r") as stats_file:
        print("PROGRESS: parsing per-sample stats", file=sys.stderr)
        for line in stats_file:
            line = line.strip().split(separator)
            sample, cloud, mass, seed, seed_abundance = line[0:5]
            index = int(seed[0:2], 16)
            if seed in per_sample_stats[index]:
                per_sample_stats[index][seed] += 1
            else:
                per_sample_stats[index][seed] = 1
            if sample != previous_sample:
                previous_sample = sample
                number_of_samples += 1

    # keep only local seeds present in at least "percentage" of samples
    threshold = percentage * number_of_samples
    seeds = [{k: v for k, v in per_sample_stats[i].items() if v >= threshold}
             for i in range(0, 256)]

    return threshold, seeds


def stats_parse(global_stats_file, threshold, seeds):
    """
    Find and eliminate global seeds.
    """
    separator = "\t"
    # 1 - a secondary seed cannot be more abundant than the global
    # seed, otherwise it would have been selected to be the global
    # seed.
    # 2 - a secondary seed cannot have less reads than the threshold
    # value, otherwise it would appear in "percentage" samples with
    # less than 1 read per sample which is not possible.
    # 3 - consequently, a global seed cannot have an abundance smaller
    # than the threshold value.
    with open(global_stats_file, "r") as stats_file:
        print("PROGRESS: parsing stats", file=sys.stderr)
        for line in stats_file:
            line = line.strip().split(separator)
            cloud, mass, seed, seed_abundance, singletons = line[0:5]

            # stats file is sorted by decreasing seed_abundance
            if int(seed_abundance) < threshold:
                break  # no need to read more lines (see point 3)

            # only check clusters with at least two unique sequences,
            # and with enough reads to be hosting a secondary seed
            # that passes our threshold
            if int(cloud) > 1 and int(mass) >= 2 * threshold + int(singletons):
                index = int(seed[0:2], 16)
                # eliminate global seeds from the list of seeds
                if seed in seeds[index]:
                    del seeds[index][seed]

    return None


def swarms_parse(swarms_file, seeds):
    """
    Map amplicons and abundance values. Only keep clusters with local seeds.
    """
    # Can a global seed be absent from the local seed list? Yes, it is
    # possible: an amplicon can be very abundant in one sample, and a
    # nearly-identical amplicon can be present in many samples but
    # with a lower total abundance. Hence, I can only rely on a search
    # through the "swarms" file to get a full list of clusters that
    # can be cleaved.
    separator = "_|;size=|;? "  # parsing of abundance annotations
    swarms = [dict() for i in range(0, 256)]
    global_seeds = [dict() for i in range(0, 256)]
    seeds_set = {k for d in seeds for k in d.keys()}
    number_of_seeds = len(seeds_set)

    with open(swarms_file, "r") as swarms_file:
        print("PROGRESS: parsing swarms", file=sys.stderr)
        for line in swarms_file:
            line = line.strip()
            amplicons = re.split(separator, line)[0::2]
            abundances = re.split(separator, line)[1::2]
            seed = amplicons[0]
            common = seeds_set & set(amplicons)
            if common:
                number_of_seeds -= len(common)
                for amplicon, abundance in zip(amplicons, abundances):
                    index = int(amplicon[0:2], 16)
                    swarms[index][amplicon] = int(abundance)
                for amplicon in common:
                    index = int(amplicon[0:2], 16)
                    global_seeds[index][amplicon] = seed
            if number_of_seeds == 0:
                break

    return swarms, global_seeds


def struct_parse(struct_file, seeds, global_seeds):
    """
    carve out sub-clusters.
    """
    separator = "\t"
    global_seeds_set = {v for d in global_seeds for v in d.values()}
    number_of_seeds = sum([len(d) for d in seeds])
    previous_cluster_id = 0
    clusters = dict()
    new_clusters = list()

    with open(struct_file, "r") as struct_file:
        print("PROGRESS: parsing struct", file=sys.stderr)
        for line in struct_file:
            line = line.strip()
            father, son, diffs, cluster_id, steps = line.split(separator)

            # initialize per-cluster parameters
            if int(cluster_id) != previous_cluster_id:
                if clusters:  # save previous results, if any
                    new_clusters.append(clusters)
                previous_cluster_id = int(cluster_id)
                global_seed = father
                has_a_local_seed = (True if global_seed in global_seeds_set
                                    else False)
                clusters = dict()
                if has_a_local_seed:
                    clusters[global_seed] = set([global_seed])

            # stop parsing the file as soon as possible
            if number_of_seeds == 0:
                break

            # skip clusters that do not contain local seeds
            if has_a_local_seed is False:
                continue

            # detect local seeds
            index = int(son[0:2], 16)
            if son in seeds[index]:
                number_of_seeds -= 1
                clusters[son] = set([son])
                continue

            # populate cluster (assuming a father-son link)
            for d in clusters:
                if father in clusters[d]:
                    clusters[d].add(son)
                    break
            else:
                # father is not in the sub-clusters (i.e. an "orphan"
                # created by the grafting process)
                for d in clusters:
                    if son in clusters[d]:
                        clusters[d].add(father)

    return new_clusters


def add_abundance_values(swarms_file, new_clusters, swarms):
    """
    Add abundance values and sort (deal with a rare case).
    """
    print("PROGRESS: sorting each cluster", file=sys.stderr)
    new_clusters_with_abundance = copy.deepcopy(new_clusters)  # suboptimal
    for i, super_cluster in enumerate(new_clusters):
        for cluster in super_cluster:
            swarm = list()
            for amplicon in super_cluster[cluster]:
                index = int(amplicon[0:2], 16)
                abundance = swarms[index][amplicon]
                swarm.append((amplicon, abundance))
            # sort amplicons by decreasing abundance value and by
            # name (fix a rare bug: a tie leading to wrong seed
            # selection)
            swarm.sort(key = lambda x: (-x[1], x[0]))

            # check if the seed changed after sorting (rare case)
            seed = swarm[0][0]
            if seed != cluster:
                del new_clusters_with_abundance[i][cluster]
                cluster = seed
            new_clusters_with_abundance[i][cluster] = swarm

    return new_clusters_with_abundance


def per_cluster_stats(global_stats_file, new_clusters_with_abundance):
    """
    Compute per-cluster stats.
    """
    print("PROGRESS: computing per-cluster stats", file=sys.stderr)
    new_stats = list()
    for super_cluster in new_clusters_with_abundance:
        for cluster in super_cluster:
            total_abundance = 0
            singletons = 0
            seed_abundance = super_cluster[cluster][0][1]
            for amplicon, abundance in super_cluster[cluster]:
                total_abundance += abundance
                if abundance == 1:
                    singletons += 1
            # number_of_uniques = 1  # 1. number of unique amplicons
            # total_abundance = 0    # 2. total abundance of amplicons
            # seed_label = 0         # 3. label of the initial seed
            # seed_abundance = 0     # 4. initial seed abundance
            # singletons = 0         # 5. number of amps with an abundance of 1
            # number_of_steps = 0    # 6. number of steps in the cluster
            # number of layers       # 7. columns 6 and 7 are not updated
            new_stats.append((len(super_cluster[cluster]),
                              total_abundance,
                              cluster,
                              seed_abundance,
                              singletons,
                              "0",
                              "0"))
    # sort clusters by increasing number of reads first, then in a
    # second step sort by decreasing number of unique amplicons, and
    # amplicon name (sort by amplicon name first, stable sorting will
    # preserve the order for ties)
    new_stats.sort(key=operator.itemgetter(2))
    new_stats.sort(key=operator.itemgetter(1, 0), reverse=True)
    with open(global_stats_file + "2", "w") as new_stats_file:
        for t in new_stats:
            print(*t, sep="\t", file=new_stats_file)

    return new_stats


def per_cluster_swarms(swarms_file, new_clusters_with_abundance):
    """
    Compute per-cluster swarms.
    """
    print("PROGRESS: computing per-cluster swarms", file=sys.stderr)
    with open(swarms_file + "2", "w") as new_swarms_file:
        for super_cluster in new_clusters_with_abundance:
            for cluster in super_cluster:
                print(*[t[0] + ";size=" + str(t[1]) for t in super_cluster[cluster]],
                      sep=" ", file=new_swarms_file)

    return None


def fasta_parse(fasta_file, new_stats, swarm_parameters):
    """
    Get seed sequences, update abundances.
    """
    new_representatives_file = (os.path.splitext(fasta_file)[0]
                                + "_"
                                + swarm_parameters
                                + "_representatives.fas2")
    # create a dict of target amplicons and abundances
    fasta = [dict() for i in range(0, 256)]
    for t in new_stats:
        index = int(t[2][0:2], 16)
        fasta[index][t[2]] = [t[1]]
    min_abundance = min([t[3] for t in new_stats])
    # filter the fasta file
    index = None
    separator = ";size="
    with open(fasta_file, "r") as fasta_file:
        print("PROGRESS: parsing fasta file", file=sys.stderr)
        for line in fasta_file:
            if line.startswith(">"):
                amplicon, abundance = line.strip(">;\n").split(separator)
                index = int(amplicon[0:2], 16)
                if int(abundance) < min_abundance:
                    break  # no need to read more lines
            else:
                if amplicon in fasta[index]:
                    fasta[index][amplicon].append(line.strip())

    with open(new_representatives_file, "w") as new_representatives_file:
        for t in new_stats:
            amplicon = t[2]
            index = int(amplicon[0:2], 16)
            try:
                abundance, sequence = fasta[index][amplicon]
            except ValueError:
                print(amplicon, fasta[index][amplicon], min_abundance)
                sys.exit(-1)
            print(">" + amplicon + separator + str(abundance)
                  + "\n" + sequence, sep="", file=new_representatives_file)

    return None


def main():
    """
    break clusters and output updated rep, stats and swarms files.
    """
    # capture arguments
    global_stats_file = ARGS.global_stats_file
    per_sample_stats_file = ARGS.per_sample_stats_file
    struct_file = ARGS.struct_file
    swarms_file = ARGS.swarms_file
    fasta_file = ARGS.fasta_file

    # fastidious or not?
    if "_1f." in swarms_file and "_1f." in struct_file:
        swarm_parameters = "1f"
    else:
        swarm_parameters = "1"

    # cleaving threshold (keystone parameter)
    PERCENTAGE = 0.05

    # Parse input files
    threshold, seeds = per_sample_stats_parse(per_sample_stats_file,
                                              PERCENTAGE)
    stats_parse(global_stats_file, threshold, seeds)
    swarms, global_seeds = swarms_parse(swarms_file, seeds)
    new_clusters = struct_parse(struct_file, seeds, global_seeds)

    # Add abundance values
    new_clusters_with_abundance = add_abundance_values(swarms_file,
                                                       new_clusters,
                                                       swarms)

    # Create output files (stats2, swarms2, fas2)
    new_stats = per_cluster_stats(global_stats_file,
                                  new_clusters_with_abundance)
    per_cluster_swarms(swarms_file, new_clusters_with_abundance)
    fasta_parse(fasta_file, new_stats, swarm_parameters)

    return


# *************************************************************************** #
#                                                                             #
#                                     Body                                    #
#                                                                             #
# *************************************************************************** #

if __name__ == '__main__':

    main()

sys.exit(0)
```

##### 2.1.2 merge sub superstring OTUs with larger OTUs

```
#!/usr/bin/env python3
# -*- coding: utf-8 -*-
"""
    merge super or sub-string OTUs with their source-OTUs.
"""

__author__ = "Frédéric Mahé <>"
__date__ = "2019/09/20"
__version__ = "$Revision: 1.2"

import sys
import csv
from optparse import OptionParser


# *************************************************************************** #
#                                                                             #
#                                  Functions                                  #
#                                                                             #
# *************************************************************************** #

def option_parse():
    """
    Parse arguments from command line.
    """
    parser = OptionParser(usage="usage: %prog -t filename -m filename",
                          version="%prog 1.0")

    parser.add_option("-t", "--table",
                      metavar="TABLE",
                      action="store",
                      dest="table",
                      help="set TABLE as input")

    parser.add_option("-m", "--matches",
                      metavar="MATCHES",
                      action="store",
                      dest="matches",
                      help="set MATCHES as input")

    parser.add_option("-o", "--output",
                      metavar="OUTPUT",
                      action="store",
                      dest="output_table",
                      help="set OUTPUT as output")

    (options, args) = parser.parse_args()

    return options.table, options.matches, options.output_table


def main():
    """
    merge super or sub-string OTUs with their source-OTUs.
    """
    input_table, matches, output_table = option_parse()
    # output_table = os.path.splitext(input_table)[0] + "_gene_counts.tsv"

    with open(matches, "r") as matches:
        OTU_connexions = dict()
        for match in matches:
            pupil, master = match.strip().split("\t")[-2:]
            OTU_connexions[pupil] = master

    # Could there be common OTUs? given the clustering process, I
    # don't think so, but I can systematically test for that.
    pupils = set(OTU_connexions.keys())
    masters = set(OTU_connexions.values())
    if masters.intersection(pupils):
        print("WARNING: there are OTUs common to hit and query columns",
              file=sys.stderr)
        print(masters.intersection(pupils), file=sys.stderr)
        sys.exit(-1)

    # Parse the OTU table
    metadata = set(["OTU", "total", "cloud", "amplicon", "length",
                    "abundance", "chimera", "spread", "quality", "sequence",
                    "identity", "taxonomy", "references"])
    with open(input_table, "r") as input_table:
        reader = csv.DictReader(input_table, delimiter="\t")
        fieldnames = reader.fieldnames
        sample_names = set(fieldnames) - metadata
        master_OTUs = dict()
        with open(output_table, "w") as output_table:
            writer = csv.DictWriter(output_table,
                                    fieldnames=reader.fieldnames,
                                    delimiter="\t")
            writer.writeheader()
            for row in reader:
                OTU = row["OTU"]
                if OTU not in masters and OTU not in pupils:
                    writer.writerow(row)
                else:
                    if OTU in masters:
                        # store it
                        master_OTUs[OTU] = row
                        for sample in sample_names:
                            master_OTUs[OTU][sample] = int(row[sample])
                            master_OTUs[OTU]["spread"] = int(row["spread"])
                            master_OTUs[OTU]["total"] = int(row["total"])
                            master_OTUs[OTU]["cloud"] = int(row["cloud"])
                    if OTU in pupils:
                        # who is its master?
                        master = OTU_connexions[OTU]
                        # update master's occurrences
                        for sample in sample_names:
                            master_OTUs[master][sample] += int(row[sample])
                            # update spread
                        new_spread = len([master_OTUs[master][sample]
                                          for sample in sample_names
                                          if master_OTUs[master][sample] > 0])
                        master_OTUs[master]["spread"] = new_spread
                        # update total
                        master_OTUs[master]["total"] += int(row["total"])
                        # update cloud
                        master_OTUs[master]["cloud"] += int(row["cloud"]) + 1
                        # output the updated masters
            for OTU in master_OTUs:
                writer.writerow(master_OTUs[OTU])

    return None


# *************************************************************************** #
#                                                                             #
#                                     Body                                    #
#                                                                             #
# *************************************************************************** #

if __name__ == '__main__':

    main()

sys.exit(0)
```

##### 2.1.3 stampa merge

(see stampa)

##### 2.1.4 OTU table updater

```
#!/usr/bin/env python3
# -*- coding: utf-8 -*-
"""
   update an OTU table with new taxonomic assignments
"""

__author__ = "Frédéric Mahé <>"
__date__ = "2020/01/08"
__version__ = "$Revision: 1.0"

import sys
import argparse


# *************************************************************************** #
#                                                                             #
#                                  Functions                                  #
#                                                                             #
# *************************************************************************** #

if __name__ == '__main__':
    """
    Parse arguments from command line.
    """
    parser = argparse.ArgumentParser(
        description="update an OTU table with new taxonomic assignments.")

    parser.add_argument("--old_otu_table",
                        dest="old_otu_table",
                        required=True,
                        help="old OTU table")

    parser.add_argument("--new_taxonomy",
                        dest="new_taxonomy_file",
                        required=True,
                        help="new taxonomic assignments")

    parser.add_argument("--new_otu_table",
                        dest="new_otu_table",
                        required=True,
                        help="new OTU table")

    ARGS = parser.parse_args()


def parse_taxonomy(new_taxonomy_file):
    """
    Map amplicons and taxonomic assignments.
    """
    separator = "\t"
    amplicons = [dict() for i in range(0, 256)]

    with open(new_taxonomy_file, "r") as new_taxonomy_file:
        print("PROGRESS: parsing taxonomy", file=sys.stderr)
        for line in new_taxonomy_file:
            amplicon, abundance, identity, taxonomy, references = \
                line.strip().split(separator)
            index = int(amplicon[0:2], 16)
            amplicons[index][amplicon] = (identity, taxonomy, references)

    return amplicons


def update_otu_table(old_otu_table, amplicons, new_otu_table):
    """
    Update taxonomy, identity and references.

    Column numbers are:
    1   OTU
    2   total
    3   cloud
    4   amplicon
    5   length
    6   abundance
    7   chimera
    8   spread
    9   quality
    10  sequence
    11  identity
    12  taxonomy
    13  references
    """
    separator = "\t"
    is_first_line = True
    with open(old_otu_table, "r") as old_otu_table:
        with open(new_otu_table, "w") as new_otu_table:
            print("PROGRESS: parsing and updating old OTU table",
                  file=sys.stderr)
            for line in old_otu_table:
                line = line.strip().split(separator)

                # header line is printed as-is
                if is_first_line:
                    is_first_line = False
                    print("\t".join(line), file=new_otu_table)
                    continue

                # update identity, taxonomy and references
                amplicon = line[3]
                index = int(amplicon[0:2], 16)
                line[10], line[11], line[12] = amplicons[index][amplicon]
                print("\t".join(line), file=new_otu_table)

    return None


def main():
    """
    update an OTU table with new taxonomic assignments.
    """
    # capture arguments
    old_otu_table = ARGS.old_otu_table
    new_taxonomy_file = ARGS.new_taxonomy_file
    new_otu_table = ARGS.new_otu_table

    # Parse the new taxomomic results
    amplicons = parse_taxonomy(new_taxonomy_file)

    # Parse the old OTU table and write a new one
    update_otu_table(old_otu_table, amplicons, new_otu_table)

    return None


# *************************************************************************** #
#                                                                             #
#                                     Body                                    #
#                                                                             #
# *************************************************************************** #

if __name__ == '__main__':

    main()

sys.exit(0)
```

### 3 phylogenetic placement

```
#!/bin/bash
cd ${HOME}/projects/Neotropical_Soils/data/

## ------------------------------------------------------ input data (unmasked)

COLPODEA="neotropical_soils_18S_V4_175_samples.OTU.filtered.cleaved.nosubstringOTUs.lulu.uchime.Colpodea.fas"
MSA_FASTA=="reference_alignment_V4_unmasked.fas"
MSA_PHYLIP="reference_alignment_V4_unmasked.phy"
NEWICK_TREE="reference_tree_V4_unmasked.tre"


## --------------------------------------------------------------------- PaPaRa
papara \
    -t "${NEWICK_TREE}" \
    -s "${MSA_PHYLIP}" \
    -q "${COLPODEA}" \
    -r

# output:
# papara_alignment.default
# papara_log.default
# papara_quality.default


## --------------------------------------------------------------------- epa-ng
epa-ng \
    --split reference_alignment_V4_unmasked.fas \
    papara_alignment.default

# output:
# query.fasta
# reference.fasta

raxml-ng \
    --evaluate \
    --msa reference.fasta \
    --tree "${NEWICK_TREE}" \
    --model GTR+G

# output:
# reference.fasta.raxml.bestModel
# reference.fasta.raxml.bestTree
# reference.fasta.raxml.log
# reference.fasta.raxml.rba
# reference.fasta.raxml.startTree

# Top 7 placements:
epa-ng \
    -t "${NEWICK_TREE}" \
    -s reference.fasta \
    -q query.fasta \
    --model reference.fasta.raxml.bestModel

# Top 70 placements:
epa-ng \
    --filter-max 70 \
    -t "${NEWICK_TREE}" \
    -s reference.fasta \
    -q query.fasta \
    --model reference.fasta.raxml.bestModel

# Top 700 placements:
epa-ng \
    --filter-max 700 \
    -t "${NEWICK_TREE}" \
    -s reference.fasta \
    -q query.fasta \
    --model reference.fasta.raxml.bestModel

# output:
# epa_result.jplace


## ---------------------------------------------------------------------- gappa
# heat tree
gappa \
    examine heat-tree \
    --jplace-path epa_result.jplace \
    --mass-norm absolute \
    --write-svg-tree \
    --write-newick-tree \
    --write-nexus-tree

# output:
# tree.svg
# tree.nexus
# tree.newick

# extraction
gappa \
    prepare extract \
    --jplace-path epa_result.jplace \
    --clade-list-file clades.txt \
    --fasta-path query.fasta \
    --color-tree-file extract_tree \
    --samples-out-dir samples \
    --sequences-out-dir sequences

# output: 
# extract.sh.txt
# extract_tree.svg
# results.log
# samples
# sequences

# per query assignment results
gappa \
    examine assign \
    --jplace-path epa_result.jplace \
    --taxon file clade.txt \
    --per-query-results

# output: 
# profile.tsv
# per_query.tsv

# LWR histogram
gappa \
    examine lwr \
    --jplace-path epa_result.jplace

# output:
# lwr_histogram.csv
# lwr_list.csv

# EDPL histogram
gappa \
    examine edpl \
    --jplace-path epa_result.jplace

# output:
# edpl_histogram.csv
# edpl_list.csv
```

### 4 R scripts

```
library(ggplot2)
library(tidyverse)
library(cowplot)

# Color palettes
cbbPalette <- c("#CC79A7", "#E69F00", "#56B4E9", "#009E73", "#000000", "#0072B2", "#D55E00", "#F0E442")


# PWC plot
(pwc <- ggplot(Histograms_BLASTall, aes(Placement, Similarity)) +
    geom_boxplot(aes(color = Placement),
                 outlier.alpha = 0) +
    scale_color_manual(values = cbbPalette,
                       guide = "none") +
    theme_classic() +
    labs(x = "pairwise alignment taxon assignment",
         y = "% similarity to reference database") +
    geom_jitter(
      aes(color = Placement),
      position = position_jitter(
        width = .2,
        seed = 2020)
    ))


# EPA plot
(epa <- Histograms_rerooted_lwr %>% 
    mutate(Placement = fct_relevel(Placement,
                                   "Bursariomorphida", "Colpodida",
                                   "Cyrtolophosidida", "Platyophryida",
                                   "outgroup")) %>%
    ggplot(aes(Placement, LWR)) +
    geom_boxplot(aes(color = Placement),
                 outlier.alpha = 0) +
    scale_color_manual(values = cbbPalette,
                       guide = "none") +
    theme_classic() +
    labs(x = "phylogenetic placement taxon assignment",
         y = "likelihood weight ratio for best placement") +
    geom_jitter(
      aes(color = Placement),
      position = position_jitter(
        width = .2,
        seed = 2020)
    ))


# EDPL plot
(edpl <- edpl_list_xls_modified %>% 
    mutate(Placement = fct_relevel(Placement,
                                   "Bursariomorphida", "Colpodida",
                                   "Cyrtolophosidida", "Platyophryida",
                                   "outgroup")) %>%
    ggplot(aes(x = Placement,
               y = as.numeric(EDPL))) +
    geom_boxplot(aes(color = Placement),
                 outlier.alpha = 0) +
    geom_point(aes(color = Placement),
               position = position_jitter(width = .2, seed = 1)) + 
    scale_color_manual(values = cbbPalette) +
    theme_classic() +
    theme(legend.position = "none") +
    labs(x = "phylogenetic placement taxon assignment",
         y = "expected distance between placement locations"))


# Perquery plot
(perquery <- per_query_results %>% 
    mutate(taxopath = fct_relevel(taxopath,
                                  "Bursariomorphida", "Colpodida",
                                  "Cyrtolophosidida", "Platyophryida",
                                  "outgroup")) %>%
    ggplot(aes(x = taxopath, 
               y = aLWR)) +
    geom_boxplot(aes(color = taxopath),
                 outlier.alpha = 0) +
    scale_color_manual(values = cbbPalette,
                       guide = "none") +
    theme_classic() +
    labs(x = "phylogenetic placement taxon assignment",
         y = "additive likelihood weight ratios for top seven placements") +
    geom_jitter(
      aes(color = taxopath),
      position = position_jitter(
        width = .2,
        seed = 2020)))


# Merging plots
library(cowplot)
plot_grid(pwc, epa, edpl, perquery, labels = c('(a)', '(b)', '(c)', '(d)'), label_size = 12)


# LWR distribution plot
# based on Lucas Czech's script:
# https://github.com/lczech/gappa/blob/master/scripts/plot-lwr-distribution.R


# Load theme
theme_set(theme_cowplot())
suppressMessages(library(viridis))

# Read table...
data <- read.table(file="lwr-distribution.csv", sep=",", header=TRUE)

# ... and turn it into the R long format, putting the previous column names into `Group`,
# and the values into `LWR`, taking all columns with LWR.x and the Remainder.
data_long <- gather(data, Group, LWR, `LWR.1`:`Remainder`, factor_key=TRUE)

# Plot the data.
ggplot(data_long, aes(x=Index, y=LWR, fill=Group)) +
  geom_area(alpha = .8, position = position_stack(reverse = TRUE)) +
  scale_fill_viridis(discrete = T) +
  coord_cartesian(ylim=c(0, 1)) +
  xlab("Pquery Index") +
  ylab("Accumulated LWR") +
  theme(legend.title=element_blank())
```

Date: 2021-04-25 dim. 00:00

Author: Frédéric Mahé

Created: 2021-04-26 lun. 17:40

Validate
